# Metric information in cognitive maps: Euclidean embedding of non-Euclidean environments

**DOI:** 10.1101/2023.06.09.544331

**Authors:** Tristan Baumann, Hanspeter A Mallot

## Abstract

The structure of the internal representation of surrounding space, the so-called *cognitive map*, has long been debated. A Euclidean metric map is the most straight-forward hypothesis, but human navigation has been shown to systematically deviate from the Euclidean ground truth. Vector navigation based on non-metric models can better explain the observed behavior, but also discards useful geometric properties such as fast shortcut estimation and cue integration.

Here, we propose another alternative, a Euclidean metric map that is systematically distorted to account for the observed behavior. The map is found by embedding the non-metric model, a labeled graph, into 2D Euclidean coordinates. We compared these two models using human data from Warren et al. (2017), where participants had to navigate and learn a non-Euclidean maze (i.e., with Wormholes) and perform direct shortcuts between different locations. Even though the Euclidean embedding cannot correctly represent the non-Euclidean environment, both models predicted the data equally well. We argue that the so embedded graph naturally arises from integrating the local position information into a metric framework, which makes the model more powerful and robust than the non-metric alternative. It may therefore be a better model for the human cognitive map.

## 1 Introduction

### 1.1 The cognitive map

The spatial long-term memory contains representations of places, landmarks and local views. A sequences of navigational actions connecting these representations is called a route and animals with such route knowledge are able to navigate between known places by following these routes (Collett et al., 1998; Collett and Collett, 2002; Warren, 2019; Mallot, 2024). If knowledge about many different items, places, and routes is integrated and novel routes and shortcuts can be inferred from previously learned route segments, the representation is called a map (Tolman, 1948; O’Keefe and Nadel, 1978; Gallistel, 1990; Trullier et al., 1997; Mallot, 2024). The cognitive map is thus a form of declarative memory in the sense that it characterizes “knowing what” or “knowing where” as opposed to the non-declarative “knowing how” of routes or guidance information (O’Keefe and Nadel, 1978; Squire and Knowlton, 1995).

A cognitive map is the most general form of spatial long-term memory, and it is believed that many animals, including humans, have access to this representation (Gallistel, 1990; Nadel, 2013; Warren, 2019). This is exemplified by the existence of neural correlates of position, the place cells (O’Keefe and Nadel, 1978; Rolls and O’Mara, 1995; Ekstrom et al., 2003; Yartsev and Ulanovsky, 2013), which encode the position of the animal within the current context via population activity.

The intuition of an internal map is relatively straight-forward, because it matches maps encountered in everyday life: In general, such maps may be broadly characterized by two frameworks: Euclidean metric maps and topological graphs. Euclidean metric maps, such as a bird’s eye view of a city or a satellite image, assign unique coordinates to each position that approximate the real-world geometry by preserving the metric relationships between positions. Topological graphs, such as a subway or bus chart or an instruction manual, describe states and possible actions that lead from one state to another, rather than geometry.

The metric framework (Fig. 1C) is considerably better suited to explain environments with a Euclidean geometric structure, and, based on the Kantian notion of an *a priori* assumption of absolute external space (Kant, 1781), it has often been argued that the cognitive map must likewise be Euclidean metric to capture these properties (O’Keefe and Nadel, 1978; Gallistel, 1990; McNaughton et al., 2006; Nadel, 2013). This theory is supported by the existence of grid cells in the entorhinal cortex, which are believed to encode metric path integration information (Hafting et al., 2005; McNaughton et al., 2006; Peer et al., 2021).

**Figure 1:**
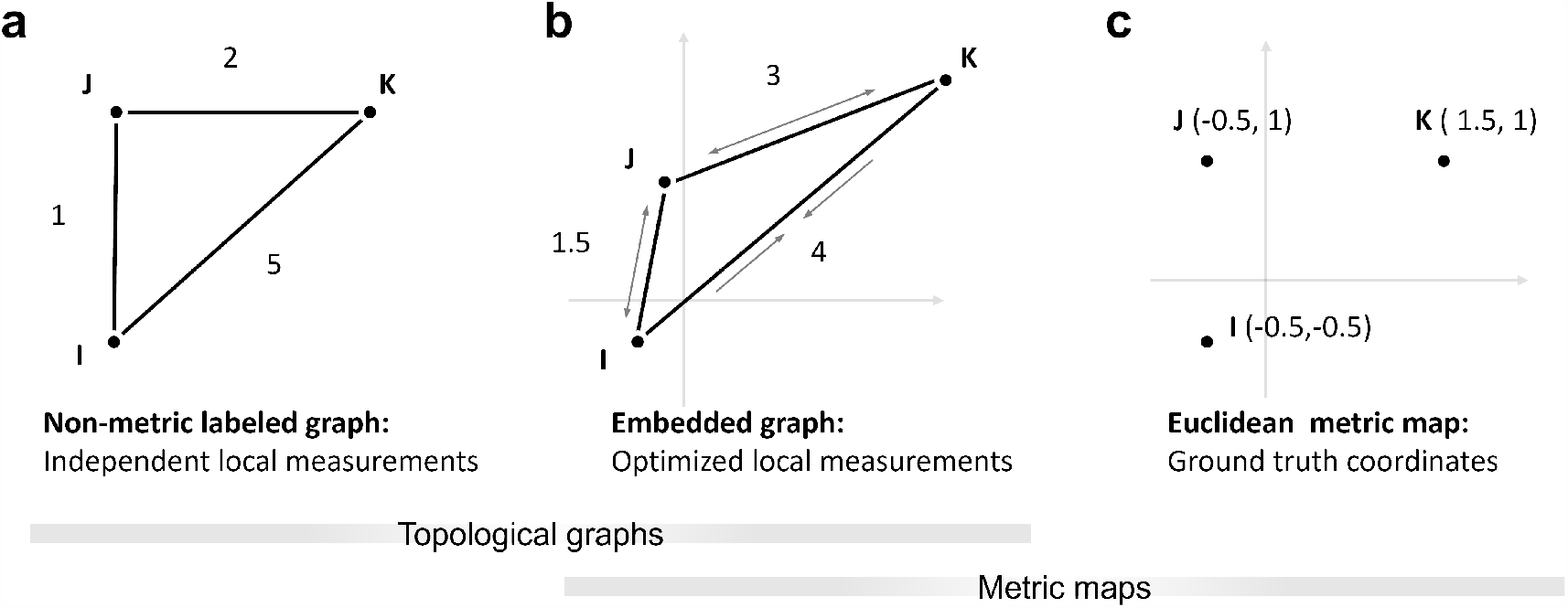
Cognitive map hypotheses. **(a)** Non-metric topological graph, labeled with distance measurements. The labels are independent of each other and do not need to adhere to the triangle inequality. **(b)** Embedded graph from (a). To find a Euclidean embedding, the distance labels need to be adjusted to create a valid configuration, for example by by stretching or compressing the edges or “wiggling” on the vertices until the difference between map and labels is minimized. As opposed to the non-metric labeled graph, changes to one label will therefore influence others. **(c)** Euclidean metric map. Places are directly assigned coordinates based on their position in the world. Over time, the coordinates may be refined by repeated measurements and the map will approach the Euclidean ground truth. The same can be expected from the embedded graph optimization if the labels are refined.

The notion of an absolute Euclidean metric may be challenged, e.g., by pointing out that the intuition of straight lines on a curved surface (or any surface that is not a plane) are actually geodesics and not true straight lines in an Euclidean sense (Helmholtz (1876); Mallot (2024), cf. Indow (1999)). But even an approximately Euclidean or non-Euclidean metric map may be advantageous, since geometric relationships between places are preserved in a highly efficient manner. That is, distances, routes, and shortcuts can be directly inferred from the map and need not be memorized individually. This property enables metric maps to store an immense amount of data, making them powerful informational tools (Nadel, 2013).

However, results from navigation experiments often disagree with the Euclidean metric map hypothesis: Human performance in shortcut or triangle completion tasks, which are often taken as evidence for an Euclidean representation, is highly unreliable with angular errors of over ±90° and angular standard deviations between 25° − 45° (Foo et al., 2005; Ishikawa and Montello, 2006; Chrastil and Warren, 2013; Warren, 2019). The Euclidean metric postulates are often violated and angle and distance estimations are systematically biased by features of the environment such as landmarks, junctions or region boundaries (Byrne, 1979; McNamara, 1986; Sadalla and Montello, 1989; Tversky, 1992; Warren, 2019; Kim and Doeller, 2022), or the number and recency of preceding turns (Brunec et al., 2017; Meilinger et al., 2018; Peer et al., 2021). In rats, place cells have been shown to stretch and shear following room deformation, while still preserving topological information about the environment (O’Keefe and Burgess, 1996; Dabaghian et al., 2014). These results imply that space is encoded much worse than what a precise metric map would predict.

As an alternative, the comparatively weaker class of topological graphs is often proposed. The environment is expressed through neighborhood or adjacency relations, forming a network of places as graph vertices and paths or actions connecting them as edges (Gillner and Mallot, 1998; Kuipers, 1978; Mallot and Basten, 2009; Warren, 2019; Peer et al., 2021). The graph may be labeled with pairwise distance or angle measurements, but this information doesn’t need to adhere to the metric postulates and is therefore not metric (Fig. 1A). Still, shortcuts and novel routes can be derived via vector addition of the labels along paths in the graph; indeed, Warren et al. (2017) suggest vector addition based on labeled graphs best explains human performance in navigation experiments. Nevertheless, poor navigational performance, biases, and large errors are not enough to completely rule out a Euclidean metric representation because the map may be systematically biased or distorted to large degrees while still being metric (Warren, 2019). Overall, the distance errors in metrically embedded maps will be smaller than in labeled graphs where distance labels are independent. Metric embedding is thus a means for efficiently exploiting all available distance information.

### 1.2 Distorted maps and non-Euclidean environments

Each individual cognitive representation will generally be different due to acquisition order, biases, and accumulation of measurement errors. One possible advantage of map-like representations in spatial memory is the mutual refinement of (possibly conflicting) local position information over time: As the agent explores the environment, it will repeatedly obtain distance and angle measurements of connections between the known places or graph states.

With a topological graph, repeated measurements of the same information could be used to create more precise labels by averaging. However, the labels will always remain independent of labels corresponding to adjacent connections and, in a triangle, might persistently violate the triangle inequality which defines a mathematical metric (Fig. 1a). In the following, we therefore refer to this representation as the *non-metric labeled graph*.

Additional precision can only be gained if repeated measurements of one connection will also improve estimates along other connections in the graph. This may for example be achieved by metric embedding (Fig. 1b). Since the acquisition of spatial memory is not complete after a single pass through the environment but relies on the consolidation of many local measurements, metric embedding seems to be a natural method for continuous integration of local information. In this sense, cue integration might be the main reason for organizing spatial representations in a metric framework.

If the measured labels are not perfect, Euclidean metric embedding can only approximate the true Euclidean metric relations and will result in a distorted depiction. The so *embedded graph* could therefore be an alternative metric explanation for the large deviations in human navigation, as opposed to the non-metric labeled graph.

In regular environments, differences between a non-metric labeled graph, an embedded graph, and a Euclidean metric map will be minimal, because the models are likely to approach the same underlying ground truth as measurements are refined. Therefore, cases need to be considered in which the models would make different predictions. With the advent of immersive virtual reality a unique opportunity has opened up to present non-Euclidean environments, thus dissociating presented metric information from the underlying true Euclidean positions (Zetzsche et al., 2009; Kluss et al., 2015; Warren et al., 2017; Widdowson and Wang, 2023). The non-Euclidean manipulations have been shown to heavily influence navigation but are usually not noticed by the subjects (Zetzsche et al., 2009; Warren et al., 2017).

### 1.3 Evidence from wormhole experiments

In the following, we focus on a specific example, Warren et al. (2017), because the experiment offers an excellent setup to investigate the hypotheses with respect to systematic distortion and the data are available online.

Warren et al. (2017) presented participants with a non-Euclidean environment and argued that, if the cognitive map is Euclidean metric, participants should have greater difficulties in learning the non-Euclidean environment compared to control, because mismatches between the cognitive map and the environment should occur. On the other hand, a non-metric graph should have no such issues.

Using head-mounted display virtual reality, Warren et al. created a hedge maze augmented with two invisible wormholes. The wormholes functioned as instant seamless teleportation and 90° rotation between different parts of the maze while participants continued to walk normally in the real-life room, therefore creating a mismatch between maze position and path integration information. Interestingly, only one participant reported noticing any kind of spatial anomaly in the maze.

Participants had to memorize object positions within the maze and were later asked to walk direct shortcuts between them. For this, the participants were moved to a starting object and had time to orient themselves. Then, the walls of the maze disappeared and the participants had to walk to the presumed position of a target object. The initial angles of the subjects’ trajectories were measured and used as directional estimates to compare the non-metric labeled graph and undistorted Euclidean map models.

Warren et al. found that directional estimates were heavily distorted towards the wormholes. This is predicted by vector addition along the shortest path on a labeled graph but not by straight lines in Euclidean ground truth coordinates. The authors thus rejected the Euclidean map hypothesis in favor of the non-metric labeled graph, arguing that only a non-Euclidean structure could explain the observed results (Warren et al., 2017; Warren, 2019). A distorted Euclidean map was briefly considered but rejected on the basis that such a map “must still satisfy the metric postulates […] in the inertial coordinate system” (Warren et al., 2017). However, the metric embedding is not a simple averaging of the path integration coordinates (Warren’s “inertial coordinates”) but the result of a optimization that also takes into account the other connections in the graph. The representation of a goal will therefore not necessarily end up in the middle between the path integration vectors obtained along two different paths, but may be closer to one or the other.

We reexamined the data used in Warren et al. (2017) with respect to the possibility of a distorted Euclidean map. In the following, we show that such a map can be found by first creating a non-metric labeled graph for the maze and then embedding the graph into 2D Euclidean coordinates. This is achieved by the minimization of the angle and distance differences between graph and map, following the method described in Hübner and Mallot (2007); Mallot (2024) for the embedding of view graphs. In an ordinary Euclidean environment, the embedding will recover the ground truth coordinates, but in a non-Euclidean environment, a residual error between embedding and local measurements must remain. Because of this error, the models should make different predictions, and may be distinguished by comparing their predictions to experimental data. That is, shortcuts derived from the embedded graph should fall somewhere between the shortcuts from the other two models.

However, we found that both models, the non-metric labeled graph and its Euclidean metric embedding, were able to predict the data equally well. Because the embedded graph is a valid Euclidean map, it is better suited for shortcut generation and especially cue integration than the non-metric alternative. We therefore refute the claim by Warren et al. (2017) that their findings cannot be explained by a Euclidean metric map, and argue for the embedded graph as a better alternative explanation.

## 2 Materials and methods

### 2.1 Data acquisition

The data used here are figures, measurements and results from Warren et al. (2017). The anonymized per-subject measurements are available as supplementary material online in the Brown University Digital Repository (http://dx.doi.org/10.7301/Z0JS9NC5, retrieved in November 2022). The relevant datasets contain measurements of the direction of individual shortcuts between object pairs in the wormhole maze, given as angular difference between the estimate and the straight line direction in Euclidean ground truth coordinates. We estimated these coordinates from pixel positions based on Fig. 2B in Warren et al. (2017) (Fig. 3A), and transformed the subject estimates into global angles (i.e., increasing counterclockwise from the positive x-axis or east). The layout of the maze and example subject estimates are shown in Fig. 2.

**Figure 2:**
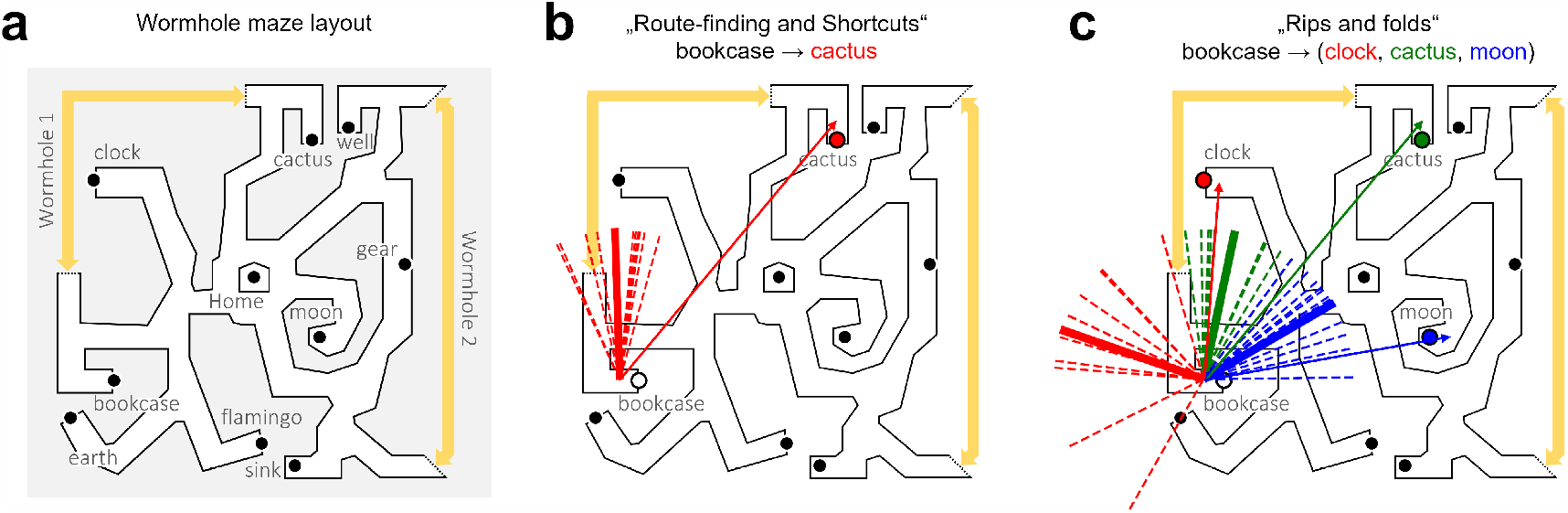
Maze and shortcut data. **(a)** Layout of the wormhole maze, redrawn from Warren et al. (2017). The yellow arrows show wormhole position and magnitude. Touching one end of the arrow instantly and seamlessly teleported subjects to the other end. **(b, c)** Example directional estimates for object pairs from the “Route-finding and shortcuts” dataset (experiment 1 in Warren et al. (2017)) and the “Rips and folds” dataset (experiment 2 in Warren et al. (2017)). The thin arrows show the Euclidean ground truth direction between objects, the short dotted lines the corresponding subject estimates, and the thick solid line the average subject estimate. The length of the estimates has been normalized and does not reflect walked distance. In (c), the colors indicate different goals.

**Figure 3:**
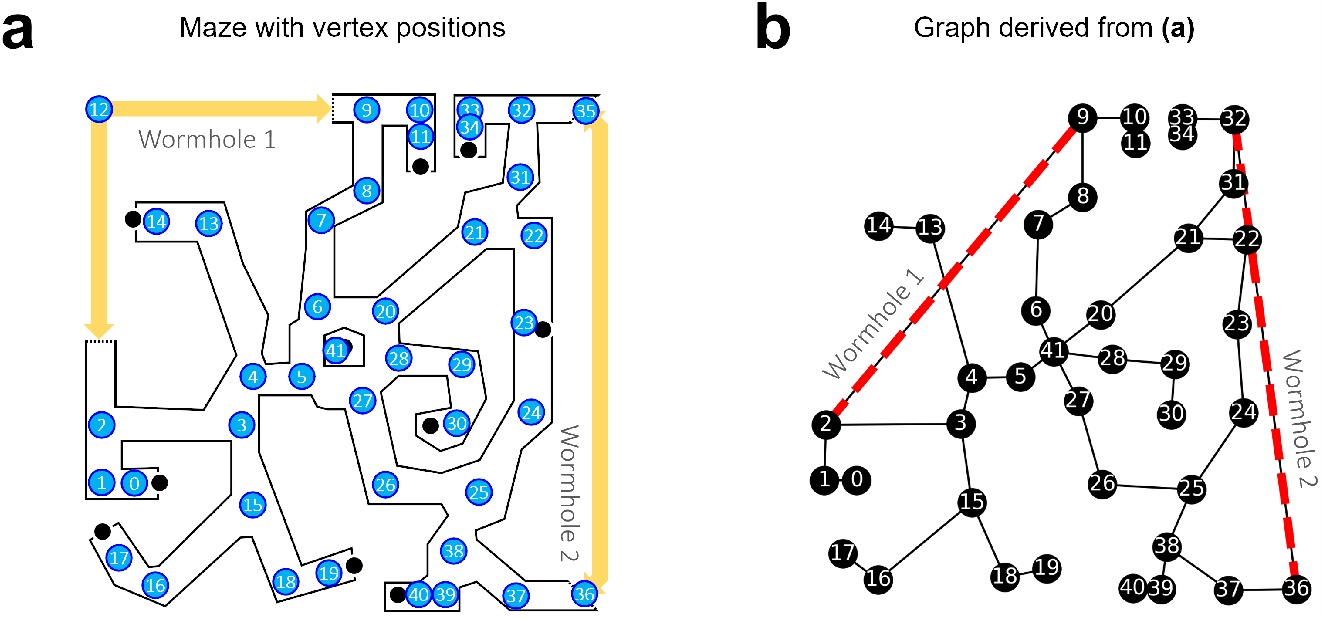
Graph creation. **(a)** Vertex positions in the maze. Their pixel coordinates were considered the Euclidean ground truth for the model. The maze was partitioned into straight segments and corners, and one vertex was placed per corner. Two vertices, 12 and 35, were only used in a control graph without wormholes. **(b)** The corresponding topological graph with edges through wormholes (red dotted lines). The graph was then labeled with local distance and angle measurements based on the ground truth, except for the wormhole edges, which were manually adjusted to reflect the locally distorted topology instead. Note that the distance along the wormhole edges is shortened but not zero.

Warren et al. (2017) measured direction estimates in two separate experiments, one to investigate shortcuts (Dataset “Route-finding and shortcuts”, see Fig. 2B) and one to investigate the ordinal reversal of landmark positions (Dataset “Rips and folds”, see Fig. 2C). “Route-finding and shortcuts” contains directional estimates of 10 subjects (5M, 5F) for four pairs of objects for a total of 10 × 4 × 2 (bidirectional) = 80 measurements. “Rips and folds” contains directional estimates of 11 subjects (9M, 2F) for eight starting locations and three targets each for a total of 11 × 8 × 3 = 264 measurements. For the purpose of this study, both datasets were treated the same but were evaluated separately; this was done for direct comparison and to avoid bias because some subjects may have participated in both studies. For further information about the participants, hardware, and experimental setup please refer to Warren et al. (2017).

### 2.2 Graph and map setup

Plausible Euclidean embeddings were found in two steps: First, a topological graph of the maze was created and labeled with the veridical distances and angles. This graph was also used to derive predictions for the non-metric labeled graph hypothesis, i.e., vector addition of the labels along the shortest paths. Next, the graph was embedded into 2D Euclidean coordinates by iterative minimization of a stress function (Hübner and Mallot, 2007; Mallot, 2024) describing the difference between the coordinates and the local labels.

In general, the creation of the topological graph is a non-trivial problem with a possibly infinite set of solutions consisting of any number of vertices, edges and measurements along the maze. Therefore, good solutions have to be guessed. Because the wormhole maze consisted of well-defined straight segments and corners, we created the graph by placing one vertex per corner and one edge per straight segment (Fig. 3B). Formally, we define the graph *G* = {*V, E*} as a set of *n* vertices *V* = {*v*_1_, …, *v*_*n*_} corresponding to places in the maze and edges *E* = {*e*_*ij*_, *e*_*jk*_, …} describing maze arms connecting the places *v*_*i*_ to *v*_*j*_ and *v*_*j*_ to *v*_*k*_.

The algorithm for metric embedding is based on local distance and turning information only, without the assumption of a global reference direction (e.g., north). It is therefore based on triplets of neighboring places *T* = {(*i, j, k*)}, i.e., places that can be visited in sequence. For each triplet, the distances *d*_*ij*_, *d*_*jk*_ and the turning angle *α*_*ijk*_ were measured and added as labels to the topological graph. *d*_*ij*_ and *d*_*jk*_ describe the distances between places *i, j* and *j, k* and *α*_*ijk*_ the heading change at *j* when moving from *i* to *k*. All labels were taken from the required egomotion steps such that labels around wormholes differed from the Euclidean ones. The same labeled graph was used for datasets from all subjects.

From the graph, a 2D Euclidean embedding *X* = {(**x**_1_, …, **x**_*n*_)} of the *n* vertices was derived by minimizing the following stress function: The algorithm considers all measured triplets of neighboring places *T* = {(*i, j, k*)} and their related distance and angle measurements (*d*_*ij*_, *d*_*jk*_, *α*_*ijk*_). Each place may appear many times as part of different triplets, and forward-backward movements of the form (*i, j, i*) are also considered (with *α*_*ijk*_ = 180°). The stress function can then be written as

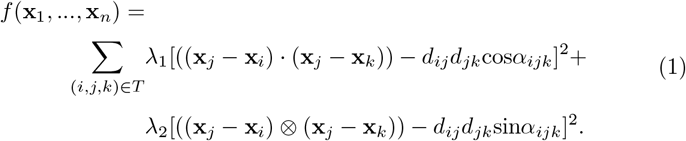

here, (·) denotes the dot product and (⊗) the third component of the cross product, (**a** ⊗ **b**) := *a*_1_*b*_2_ − *a*_2_*b*_1_, which is twice the area of the triangle (*i, j, k*). The constants *λ*_1_, *λ*_2_ can be used to weigh the components based on their variances (Mallot, 2024); we chose *λ*_1_ = *λ*_2_ = 1.

Finding an embedding that minimizes this stress function is a nonlinear optimization problem. Solutions may for example be found with iterative numerical approximations like Newton’s method. We used the quasi-Newton method Sequential Least Squares Programming (SLSQP), as implemented in the *SciPy 1*.*10 optimize* Python library (Virtanen et al., 2020), credited to (Kraft, 1988). The resulting embedding will be a Euclidean metric map of the graph’s vertices with an arbitrary global orientation, but it is not a complete distorted map of the wormhole maze in the sense that it only assigns coordinates to the vertices but not to other places. The distorted position of other places may be found by adding them as additional vertices to the graph before embedding or by interpolation. Nevertheless, the embedding is sufficient to derive directional predictions.

### 2.3 Model comparison and data analysis

Next, the non-metric labeled graph and its Euclidean embedding were used to derive predictions about shortcut directions between object pairs. For the non-metric labeled graph, predictions were obtained by finding the shortest path between start and target object using Dijkstra’s algorithm, as implemented in the *NetworkX 3*.*0* Python library (Hagberg et al., 2008). Along the path, the angles and distances were summed up to a vector, and the global direction of the resultant vector relative to the ground truth coordinates was considered the final shortcut prediction. In the embedded graph, shortcuts were simply the straight lines from start to target objects.

The predictions of the two graph models were compared to the subject data and the prediction error was measured. Because the embedded graph has no defined reference direction, subject estimates had to be considered relative to a local reference. We used the respective local angle between the starting arm and measurement or prediction, which is reference direction-independent.

For each model, the mean prediction errors and between-subject angular deviation were calculated for the group, and the within-subject angular deviation separately for each participant. The errors were compared with the two-sample Watson-Williams F-test for circular data (Batschelet, 1981), as implemented in the *PyCircStat* Python library (Berens and Sinz, 2022). The null hypothesis assumes that the samples come from underlying distributions with the same mean (Batschelet, 1981), i.e., that the models explain the subject data equally well; note that this does not mean that the models make the same prediction. Cohen’s *d* was used as a measure for effect size. All statistical tests were two-tailed with *α* = 0.05.

## 3 Results

### 3.1 Embeddings

The numerical optimization method may find different local minima. Which solution is found depends on the starting point in the solution space, i.e., the initial vertex positions *X* = {(**x**_1_, …, **x**_*n*_)}. We restarted the optimization procedure 1000 times with random initial vertex positions *X* ∼ 𝒰_2_(0, 20) and found two local minima with stress values *f* (*X*_1_) = 450.68 (Fig. 4c) and *f* (*X*_2_) = 367.36 (Fig. 4b). In the following, we report results from the first embedding, which resulted in better fits to the subject data.

**Figure 4:**
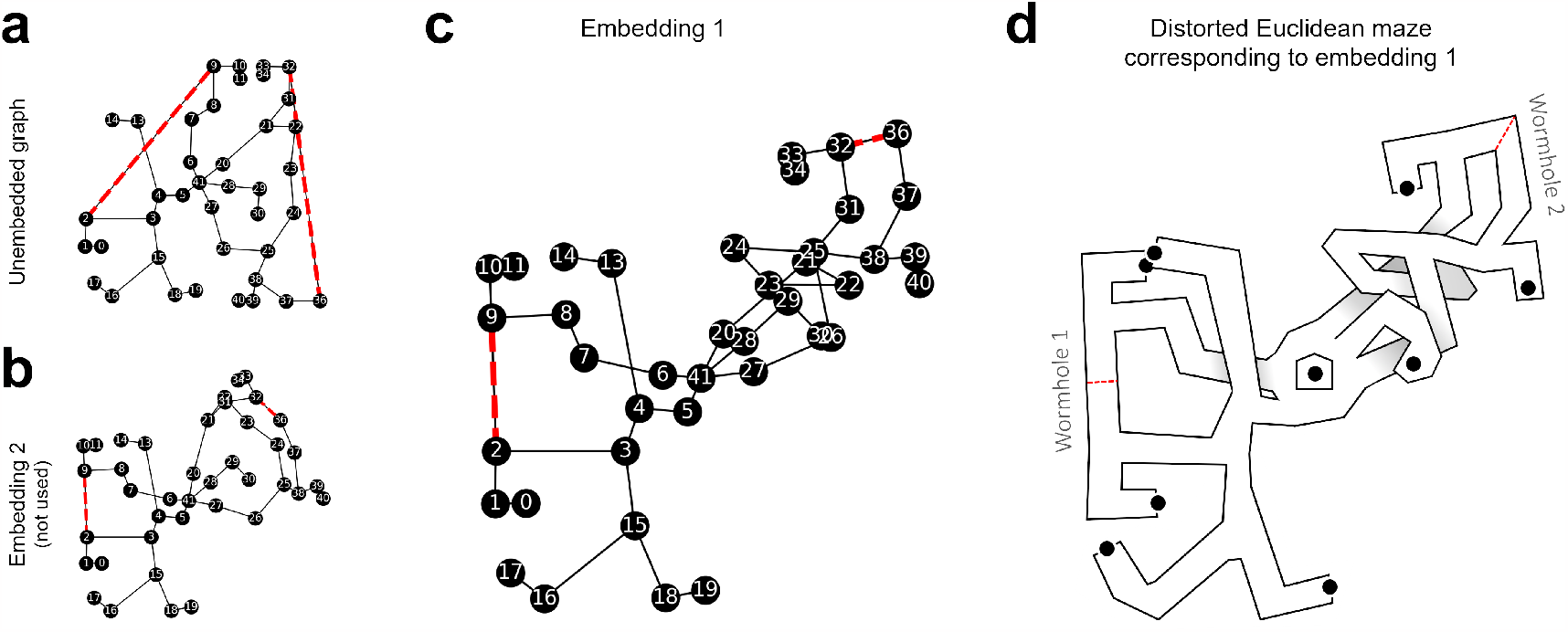
Embedded graph. **(a, b)** References for comparison. The unembedded graph (as in Fig. 3) and an another embedding which was also found by the optimization method. The second embedding performed worse on the subject data and was not further used. **(b)** The embedded graph, i.e., the labeled graph with the vertices at coordinates that minimize the difference between map and labels. The orientation of the embeddings is arbitrary; here, they were rotated so that the edge (2, 3) is horizontal. The red dotted lines show the edges that pass through wormholes. **(d)** Sketch of the distorted wormhole maze according to the embedding in (c). Edges that cross each other in the embedded graph could for example be realized through multi-level paths.

### 3.2 Dataset 1: Route-finding and shortcuts

We derived shortcut predictions from the non-metric labeled and embedded graph models and compared the predictions to human shortcut estimates from Warren et al. (2017), dataset “Route-finding and shortcuts” (Fig. 5A-C). The resulting angular prediction error was measured. Rayleigh tests on error direction revealed non-uniform distributions, *z*(10) = 9.59, *p <* .001 for the non-metric labeled graph and *z*(10) = 9.57, *p <* .001 for the embedding.

**Figure 5:**
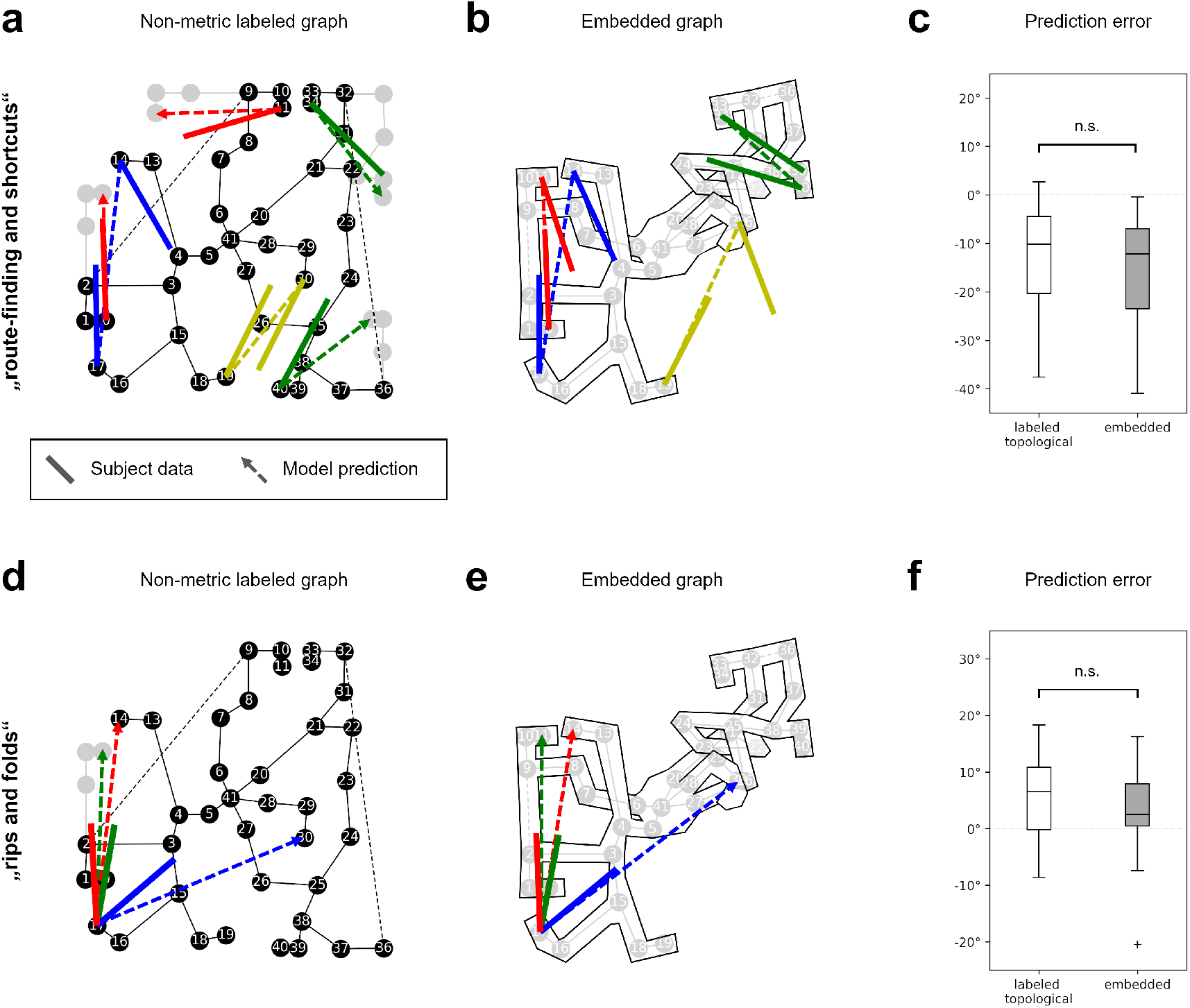
Results. (a-c): Dataset “route-finding and shortcuts”, (d-f): Dataset “rips and folds”. **(a)** Shortcut predictions of the non-metric labeled graph (dotted lines) and average subject estimates (solid lines), plotted on ground truth coordinates. The gray vertices show how the graph would continue on routes through wormholes. **(b)** Shortcut predictions of the embedded graph, lines as in (a). The subject estimates were rotated to match the local orientation of the originating maze arm. **(c)** Distribution of the prediction error. The difference between the models is not significant, i.e., they predict the data equally well. **(d)** Example shortcut predictions (dotted lines) and subject estimates (solid lines) for three of the 24 object pairs in the “rips and folds” dataset. **(e)** Shortcut predictions of the embedded graph for the same object pairs as in (d). **(f)** Distribution of the prediction error. The difference between the models is also not significant on this dataset.

The non-metric labeled graph model showed an average angular error of −12.4° with an angular deviation (*AD*) of 11.76° and the embedding an average error of −15.26°, *AD* = 11.98°. This difference was not significant (*F* (1, 18) = 0.2, *p* = .63) with a small effect size (*d* = .22). I.e., the shortcut directions derived from the graph model were not significantly closer to the subject data than the shortcut directions derived from the embedding or vice versa.

The within-subject angular deviation of the errors was fairly high but also similar for both models, with an average of 29.75° for the graph model and 32.15° for the embedding. Statistical comparison (*F* (1, 18) = 0.6, *p* = .42, *d* = .51) again revealed no significant difference.

### 3.3 Dataset 2: Rips and folds

For the purpose of this study, the “Rips and folds” dataset was treated the same as the “Route-finding and shortcuts” dataset, with the only difference being the number of participants (11 vs. 10 in dataset 1) and estimates per participant (24 vs. 8 in dataset 1). The datasets were analyzed separately for the sake of comparison.

We again compared prediction errors of the non-metric labeled graph model and its Euclidean embedding (Fig. 5D–F). Rayleigh test on error direction revealed non-uniform distributions, *z*(11) = 10.78, *p <* .001 for the non-metric labeled graph and *z*(11) = 10.77, *p <* .001 for the embedded graph. The non-metric labeled graph showed an average angular error of 5.68°, *AD* = 8.12°, and the embedded graph an error of 2.37°, *AD* = 9.35°. This difference was again not significant (*F* (1, 20) = 0.72, *p* = .41) with a small effect size (*d* = .39). Within-subject angular deviation of the errors was also high, with an average of 42.36° for the non-metric labeled graph model and 33.77° for the embedding. This difference was trending towards significance (*F* (1, 20) = 4.07, *p* = .057) with a large effect (*d* = .85).

Although dataset 2 contained many more measurements than dataset 1, there was still no significant difference between the prediction errors, i.e., the models again predicted the data equally well, with the embedding possibly capturing the within-subject variation better.

## 4 Discussion

Using subject data from Warren et al. (2017), we compared two cognitive map models, the non-metric labeled graph and the embedded graph. We found both models predicted the data equally well, i.e., both models made prediction errors with a similar magnitude and distribution. The embedding may possibly be somewhat better at predicting the within-subject angular deviation in the rips and folds dataset, but the results did not pass the selected significance threshold at *α* = .05. Given the data, we therefore found insufficient evidence to reject the null hypothesis.

Due to the non-Euclidean property of the environment, a perfect Euclidean embedding does not exist and a difference between the models must remain. It is therefore surprising that it did not lead to significantly different prediction errors.

In the original study, subjects explored the environment by walking continuous paths and thereby obtained information not only about the place-to-place distances and turns but also about the overall connectivity of the network. The conclusions drawn in Warren et al. (2017) imply that this network information is not used for the shortcut task, which is thought to be solved by vector addition along the direct path only. Here, we showed that the behavioral data are also consistent with the idea of consolidating both distance and network information in a metrically embedded graph. We thus refute the conclusion in Warren et al. (2017): it is not necessary to discard Euclidean metric properties and to reduce the representation to a non-metric framework in order to explain the observed behavior.

The main difference between the vector navigation in the labeled graph and the embedded graph suggested here lies in the treatment of repeated distance and angle measurements during prolonged navigation. Repeated measurements might simply be used to improve the estimates of distances and angles for individual labels without exploiting the constraints that these measurements impose on adjacent labels and indeed on the entire graph. Metric embedding, in contrast, allows to make use of these constraints such that improved estimates of one edge will lead to better distance and angle estimates everywhere. In this view, the main advantage of having a metrically embedded representation of space is not so much its resemblance to a geographic map, but the possibility to integrate local and repeated measurements into a consolidated structure. The result is still a graph, but with metrically embedded vertices from which directions can be derived directly without the “mental path integration” procedure suggested by Warren et al. (2017).

Note that even an optimal metric embedding is not necessarily equivalent to the Euclidean ground truth; cognitive space is not natural space and the internal representation may still be systematically distorted, even under normal Euclidean circumstances. This might explain poor navigational performance even after prolonged exposure to the environment (e.g., Ishikawa and Montello (2006)).

Non-metric topological and metrically embedded information may also coexist. Combined models have previously been proposed, for example for different levels of spatial hierarchy (Couclelis et al., 1987; Meilinger, 2008), where the local Euclidean structure of individual places or regions is known but higher-level relations between different regions are encoded as a graph. For example, a local plaza may be well-represented by a Euclidean metric map, but directions to other places within the city may only be memorized as a sequence of turns. In the context of this present study, this relates to the problem of what constitutes a vertex of the graph. In our simulation, we placed vertices at all corners of the maze, but other choices are possible. A neural network model assuming metric representations within small regions and categorical knowledge of these regions themselves has been presented by Baumann and Mallot (2023).

Topological and metric information may also be used under different environmental constraints or at different stages of exploration and familiarization (Peer et al., 2021). Initially, the environment may be encoded in terms of adjacency relations and individual routes, which then over time is consolidated in an encompassing map as the amount of information increases. This scenario is supported by reports that grid cell firing fields are initially anchored by the walls of individual compartments, but with experience extend across boundaries to encompass a larger space (Carpenter et al., 2015; Wernle et al., 2018). The embedding algorithm presented here may also be considered a support, because it describes a transformation of local position information under topological constraints into a Euclidean metric map.

## Compliance with Ethical Standards

### Data availability statement

Data will be made available upon request.

### Author contribution

**TB** Conceptualization, Formal analysis, Software, Writing; **HAM** Conceptualization, Writing.

### Funding information

The research reported in this paper was carried out at the Department of Biology of the University of Tübingen. This research did not receive any specific grant from funding agencies in the public, commercial, or non-profit sectors.

